# Single-dose YF17D-vectored Ebola vaccine candidate protects mice against both lethal surrogate Ebola and yellow fever virus challenge

**DOI:** 10.1101/2023.01.13.523888

**Authors:** Viktor Lemmens, Lara Kelchtermans, Sarah Debaveye, Winston Chiu, Thomas Vercruysse, Ji Ma, Hendrik Jan Thibaut, Johan Neyts, Lorena Sanchez-Felipe, Kai Dallmeier

## Abstract

Ebola virus (EBOV) and related filoviruses such as Sudan virus (SUDV) threaten global public health. Effective filovirus vaccines are available only for EBOV, yet restricted to emergency use considering a high reactogenicity and demanding logistics. Here we present YF-EBO, a live YF17D-vectored dual-target vaccine candidate expressing EBOV-glycoprotein (GP) as protective antigen. Safety of YF-EBO in mice was further improved over that of parental YF17D vaccine. A single dose of YF-EBO was sufficient to induce high levels of EBOV-GP specific antibodies and cellular immune responses, that protected against lethal infection using EBOV-GP-pseudotyped recombinant vesicular stomatitis virus (rVSV-EBOV) in interferon-deficient (*Ifnar*^*-/-*^) mice as surrogate challenge model. Concomitantly induced yellow fever virus (YFV)-specific immunity protected *Ifnar*^*-/-*^ mice against intracranial YFV challenge. YF-EBO could thus help to simultaneously combat both EBOV and YFV epidemics. Finally, we demonstrate how to target other highly pathogenic filoviruses such as SUDV at the root of a current outbreak in Uganda.

## Introduction

Ebola virus (EBOV), a member of the *Filoviridae* family, is causing a severe and acute systemic disease in humans known as Ebola virus disease (EVD) with mortality rates up to 80%^1^. Whereas after its discovery in 1976, only sporadic local outbreaks were reported on the African continent, more recently EVD is on the rise sparking large epidemics in West Africa (2013-16) and the Democratic republic of Congo (DRC) (2018-2020, and 2021), deteriorating societies, economies and political systems in already unstable regions.

The devastating 2013-2016 West-African epidemic – with more than 28.000 confirmed cases, 11.000 deaths and an estimated economic cost of over USD 50 billion^2^ – fostered an accelerated development of EBOV vaccines to at least protect frontline healthcare workers, of which two have been granted conditional licensure by the European Medicines Agency. Ervebo® (rVSV-EBOV) that is based on replication-competent recombinant vesicular stomatitis virus (VSV) expressing the EBOV glycoprotein (GP) proved effective in an outbreak response^3,4^. However, its high reactogenicity with fever, arthralgia, fatigue and viral dissemination in joints and skin ^5–7^, next to a strict requirement for ultradeep cooling during storage and transport precludes its wider use in routine immunization. Likewise, for Zabdeno/Mvabea® (Ad26-ZEBOV/MVA-BN-Filo) a lower estimated efficacy^8^ and an extended 3-month prime-boost regimen are disadvantages limiting its use in the field ^9^. Hence potent, safe and convenient second-generation vaccines need to be developed to protect risk populations in EBOV-endemic regions^10,11^.

Yellow fever virus (YFV) is a mosquito-borne flavivirus causing severe hemorrhagic disease in humans. Yellow fever (YF) is endemic in Central and South America, as well as sub-Saharan Africa, where EVD surges. Despite the availability of a very efficient live-attenuated yellow fever vaccine (YF17D), annually an estimated 51,000–380,000 severe cases of YF still occur resulting in 19,000–180,000 deaths^12^. Re-emergence of YF outbreaks can be mainly attributed to a low vaccine coverage due to supply issues of the vaccine^13^. Therefore, alike for EVD, second-generation YFV vaccines with a sustainable supply are needed^14,15^.

YF17D is considered one of the most effective vaccines, stimulating a broad range of innate immune responses resulting in strong humoral and polyfunctional cellular immune responses, which can provide long lasting protection after a single dose^16^. Due to its excellent immunogenic properties, YF17D has been successfully used as a vector platform for the development of novel vaccines^17^.

Since YFV and EBOV share their endemicity, a single vaccine that could simultaneously contribute to the elimination of yellow fever epidemics and combat surges of EVD would be of great benefit in routine immunization programs in endemic regions. Here we report on the development of such a dual-target YF17D-vectored Ebola vaccine candidate (YF-EBO) expressing EBOV-GP from a replication-competent full-length YF17D backbone. A single shot of YF-EBO induces both EBOV- and YFV-specific humoral and cellular immune responses in mice. This translated into dual protection from both stringent surrogate EBOV and YFV challenge in IFN type I receptor-knockout (*Ifnar*^*-/-*^) mice.

## Results

### Generation and characterization of YF-EBO

YF-EBO was generated by inserting the sequence of the EBOV (Makona strain) glycoprotein (EBOV-GP) as translational in-frame fusion at the E/NS1 intergenic region of an infectious cDNA clone of YF17D^17,18^ (**Fig. 1A**). The YF-EBO vaccine construct was rescued by transfection into BHK-21J cells, which yielded infectious viral progeny. Consistent with the replicative trade-off posed by the insertion of foreign genes into the YF17D backbone, YF-EBO had a smaller plaque phenotype (**Fig. 1B**) and showed somewhat reduced viral growth kinetics on BHK-21J cells as compared to parental YF17D (**Fig. 1C**). Expression of both YF17D and EBOV-GP antigens was confirmed by immunofluorescent staining of YF-EBO-infected BHK-21J cells with polyclonal YF17D antiserum and an EBOV-GP specific antibody (**Fig. 1D**)

**Figure 1:**
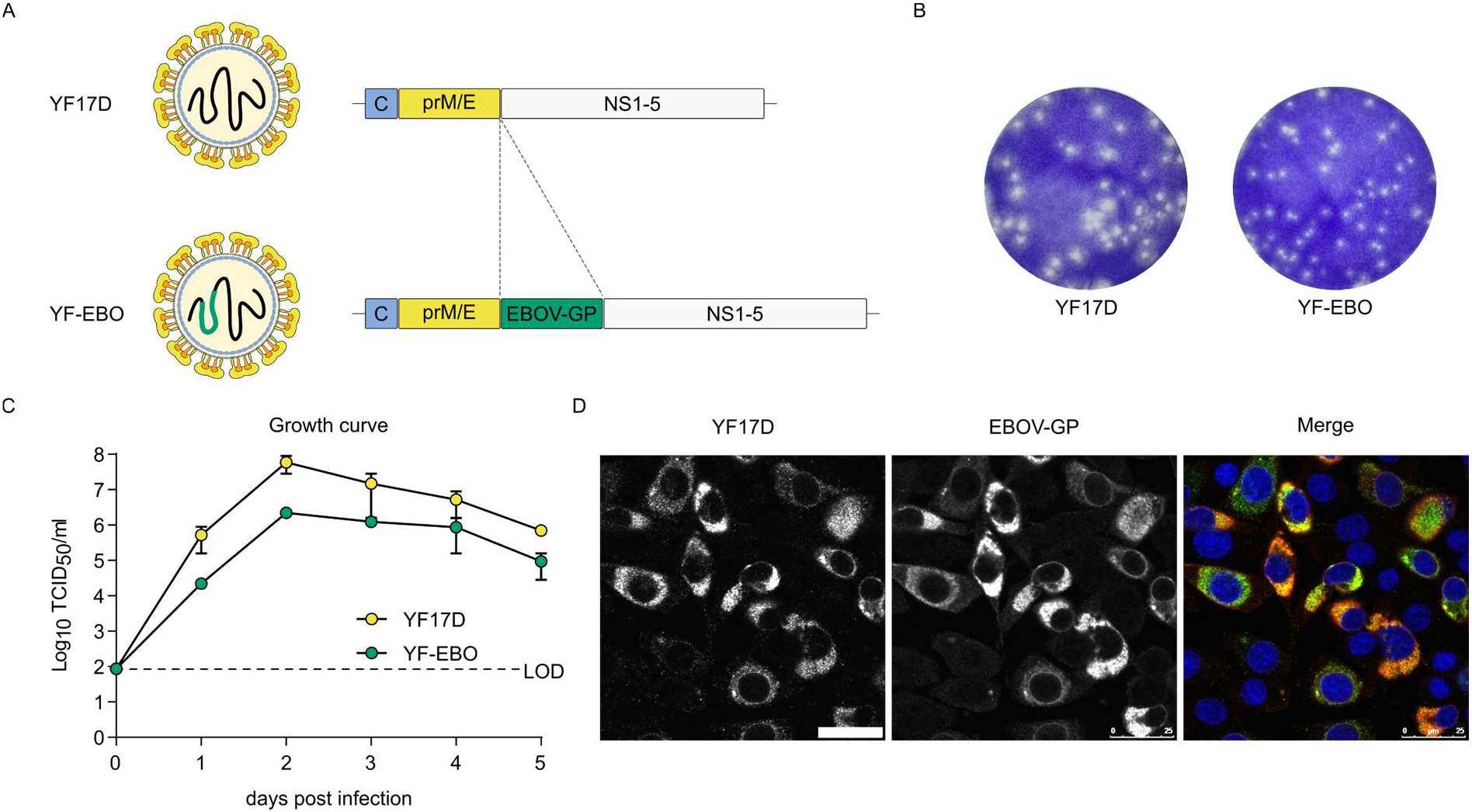
vaccine design and *in vitro* characteristics. **A**. Schematic of YF17D and YF17D-vectored Ebola vaccine candidate (YF-EBO). **B**. Representative images of plaque phenotypes from YF17D and YF-EBO on BHK-21J cells, fixed six days post-infection. **C**. Growth kinetics of YF17D and YF-EBO. BHK-21J cells were infected at a multiplicity of infection (MOI) of 0.01 and virus yields were quantified over time by virus titration on BHK-21J cells. Error bars indicate SEM (*n* = 2) and dashed line represents limit of detection (LOD). **D**. Antigenicity of YF-EBO: confocal immunofluorescent images of BHK-21J cells two days post-infection with YF-EBO, staining for YF17D (green) and EBOV-GP antigen (red) (nuclei stained with DAPI, blue). Scale bar, 25 μm.

The safety and attenuation of YF-EBO was compared to YF17D using different mouse models (**Fig. 2**). First, intraperitoneal inoculation of YF-EBO into six-to eight-week-old IFN type I receptor-knockout mice (*Ifnar*^*-/-*^) was well tolerated, as concluded from a normal weight gain over time (**Fig. 2A**). To evaluate the neuroinvasive properties of YF-EBO we used IFN type I and II receptor-knockout mice (AG129) which are highly susceptible to neuroinvasive YF17D infection^18^. Intraperitoneal inoculation of 250 plaque forming units (PFU) of YF-EBO (*n* = 8) did not result in any disease symptoms in AG129 mice, whereas a similar dose of YF17D induced neurological symptoms (e.g., paresis, paralysis or a hunched back), leading to lethality in the vast majority of mice (*n* = 5 of 6) (**Fig. 2B**). Finally, a neurovirulence test resembling a classical YF17D neuropotency test in BALB/c pups was performed by inoculating five-day-old pups intracranially with 25 PFU of YF-EBO (*n* = 12) or 10 PFU of YF17D (*n* = 10). Even when using this higher dose of YF-EBO, a slight improvement regarding survival was observed compared to YF17D inoculation (**Fig. 2C**) (median time to humane endpoint of 9 days, interquartile range (IQR) of 8.25-9 versus 8 days, IQR 8-8, respectively; *p* <0.0001, log-rank-test).

**Figure 2:**
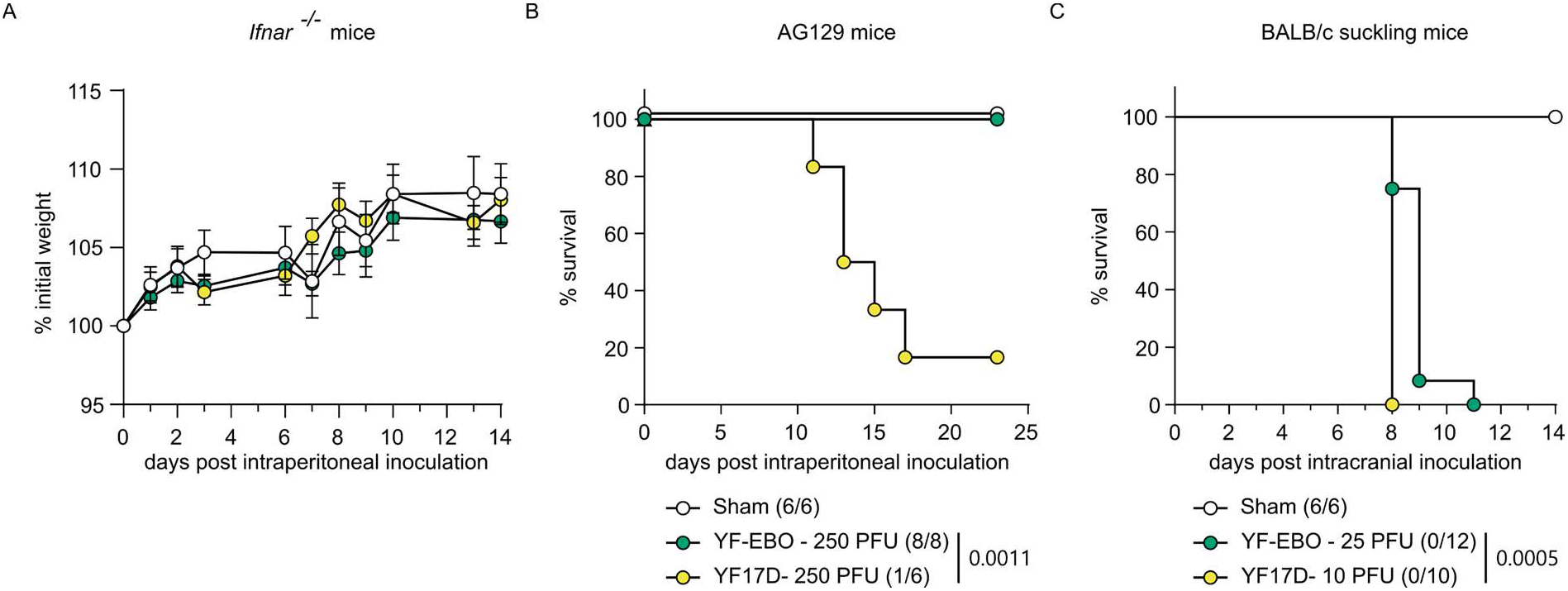
attenuation of YF-EBO. **A**. Weight evolution of *Ifnar* ^*-/-*^ mice after intraperitoneal inoculation with 250 PFU of YF-EBO (*n* = 6, green circles), YF17D (*n* = 6, yellow circles), or sham (*n* = 6, white circles) as a control. Error bars indicate SEM. **B**. Survival curve of AG129 mice after intraperitoneal inoculation with 250 PFU of YF-EBO (*n* = 8), YF17D (*n* = 6) or sham (*n* = 6) as a control. **C**. Survival curve of five-day-old suckling BALB/c mice after intracranial inoculation with 25 PFU of YF-EBO (*n* = 12), 10 PFU of YF17D (*n* = 10) or sham (*n* = 6) as a control. The number of surviving mice at study endpoint are indicated within parentheses in the legends and a log-rank test was applied to compare YF-EBO-with YF17D-vaccinated mice, significant *p*-values < 0.05 are indicated (B, C).

### EBOV-specific immunogenicity and protection against lethal rVSV-EBOV infection in mice

We used rVSV-EBOV, a replication-competent recombinant VSV pseudotyped with EBOV-GP resembling commercial Ervebo®, as a surrogate for EBOV infection in *Ifnar*^*-/-*^ mice. Likewise, *Ifnar*^*-/-*^ mice are susceptible to YF17D vaccination and hence widely used as model to study YF17D and YF17D-vectored vaccines^19^. rVSV-EBOV has previously been shown to cause lethal infection in *Stat2*^*-/-*^ mice^20^, which alike *Ifnar*^*-/-*^ mice lack a functional antiviral interferon (IFN) type I response. Importantly, rVSV-EBOV is a biosafety level (BSL)-2 agent, thus not requiring BSL-4 containment as original EBOV.

To establish our rVSV-EBOV challenge model, *Ifnar*^*-/-*^ mice were infected intraperitoneally with either 100 (*n* = 3) or 100.000 (*n* = 3) plaque forming units (PFU) of rVSV-EBOV and monitored for the development of disease. A rapid onset of disease symptoms such as weight loss (**Fig. S1A**), inactivity, ruffled fur, hunched posture and lethargy was observed for both doses, resulting in the development of high pain scores (**Fig. S1B**) ultimately requiring euthanasia (**Fig. S1C**). Organs (liver, spleen, kidney, brain and lung) were collected at the day of euthanasia to determine infectious viral loads and rVSV-EBOV replication was found in all collected organs (**Fig. S1D**). The highest infectious viral loads, up to 10^8^ TCID_50_ /30mg organ, were detected in liver and spleen. Thus, similar to *Stat2*^*-/-*^ mice, we observed that intraperitoneal inoculation of only 100 PFU of rVSV-EBOV in *Ifnar*^*-/-*^ mice results in a systemic lethal infection with rapid onset of disease symptoms. Latter model was used in the following to assess vaccine efficacy of YF-EBO.

To characterize the EBOV-specific immune responses induced by YF-EBO, we vaccinated six-to eight-week-old *Ifnar*^*-/-*^ mice with a single low dose of 250 PFU^19^ of either YF-EBO, YF17D as a matched placebo or sham as a negative control (**Fig. 3A**).

**Figure 3:**
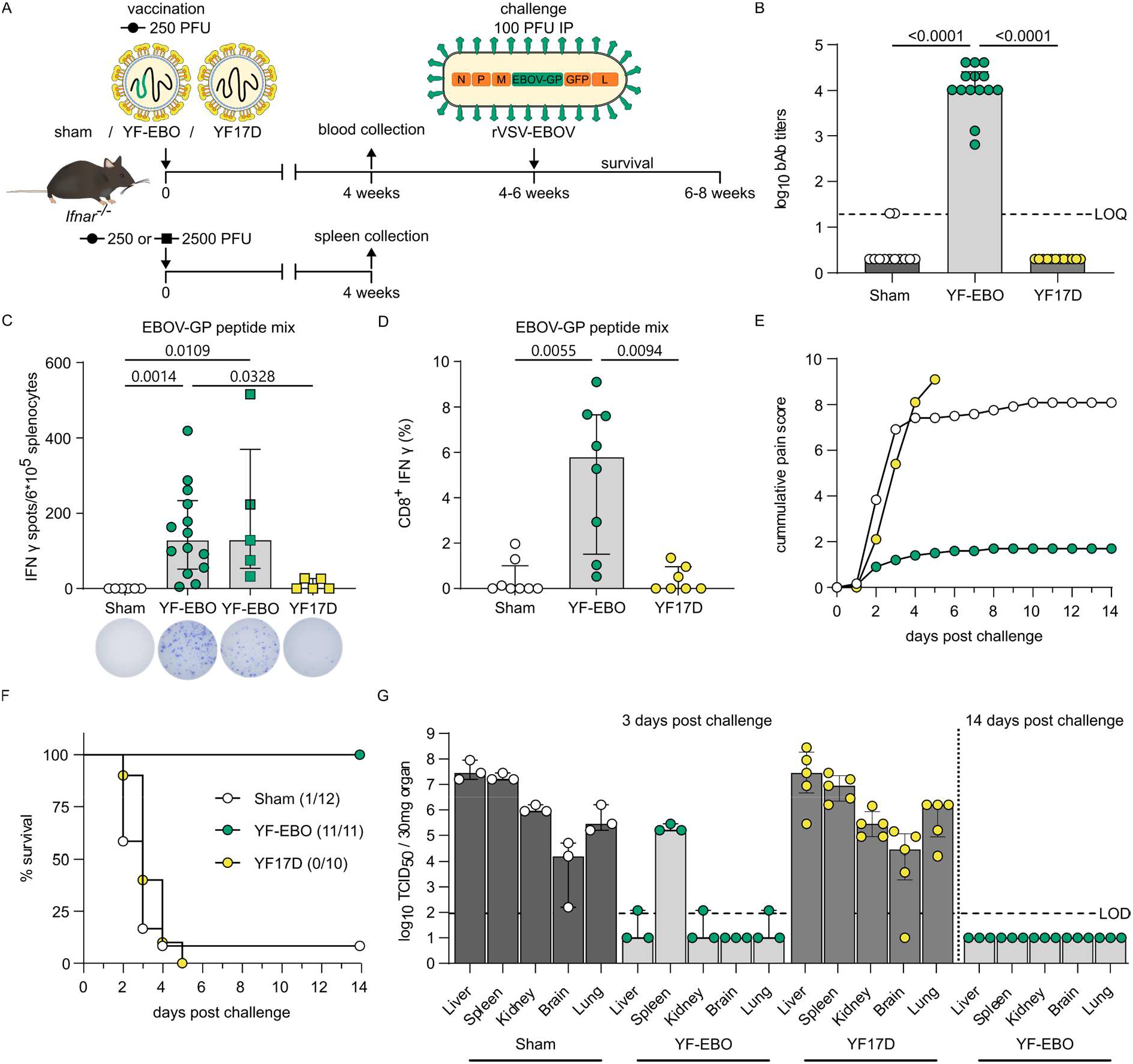
EBOV-specific immunity and protection in mice. **A**. *Ifnar* ^*-/-*^ mice were vaccinated intraperitoneally with 250 (circles) or 2500 PFU (squares) of either YF-EBO, YF17D or sham. A subset of mice vaccinated with 250 PFU and all mice vaccinated with 2500 PFU were sacrificed four weeks post-vaccination for spleen collection. Remaining mice vaccinated with 250 PFU were challenged intraperitoneally with 100 PFU of rVSV-EBOV four to six weeks post-vaccination after which they were monitored daily for two weeks for the development of disease symptoms. **B**. Pre-challenge EBOV-GP-specific IgG binding antibody (bAb) titers determined by IIFA at four weeks post-vaccination with 250 PFU (YF-EBO *n* = 14; YF17D *n* = 12; sham *n* = 12). **C**. ELISpot counts of IFNγ-secreting cells after EBOV-GP peptide pool stimulation of isolated splenocytes from mice vaccinated with 250 PFU (circles; YF-EBO *n* = 14; sham *n* = 6) or 2500 PFU (squares; YF-EBO *n* = 5; YF17D *n* = 5). A representative image of an ELISpot well from each group is shown below the x-axis. **D**. Percentage of IFNγ-expressing CD8^+^ cells after EBOV-GP peptide pool stimulation of splenocytes from mice vaccinated with 250 PFU (YF-EBO *n* = 8; YF17D *n* = 7; sham *n* = 8). **E**. Mean cumulative pain scores of mice post-challenge (YF-EBO *n* = 11; YF17D *n* = 10; sham *n* = 12), determined based on IACUC parameters including: body weight changes, body condition score, behaviour and physical appearance. **F**. Percentage survival post-challenge, the number of surviving mice at study endpoint are indicated within parentheses. **G**. rVSV-EBOV infectious viral loads in different organs at 3 days (YF-EBO *n* = 3; YF17D *n* = 4; sham *n* = 3) and 14 days post-challenge (YF-EBO *n* = 3) quantified by virus titration on Vero E6 cells. Dashed line indicates limit of quantification (LOQ) or limit of detection (LOD). Data are median ±IQR (B-D and G) and two-tailed Kruskal–Wallis test was applied followed by Dunn’s multiple comparison, significant *p*-values < 0.05 are indicated (B-D).

Four weeks post-vaccination, all YF-EBO-vaccinated mice (*n* = 14 of 14) had seroconverted to high levels of binding antibodies (bAb) as determined by indirect immunofluorescence assay (IIFA) for EBOV-GP-specific IgG (**Fig. 3B**). As for other EBOV vaccines^21,22^, EBOV-GP-specific neutralizing antibodies (nAbs) could not readily be detected (**Fig. S2A**).

Cell mediated immune (CMI) responses were analyzed four weeks post-vaccination in splenocytes isolated from a subset of vaccinated mice. Splenocytes of YF-EBO-vaccinated mice with a single low dose of 250 PFU (*n* = 14) had a significantly higher number of IFNγ-secreting virus-specific T lymphocytes measured by ELISpot after stimulation with an EBOV-GP peptide mix as recall antigen suggestive for successful induction of an antiviral T helper 1 (T_h_1)-polarization of CMI responses (**Fig. 3C**) as compared to sham (*n* = 6; *p* = 0,0014 Kruskal-Wallis-test). A higher dose of 2500 PFU of YF-EBO (*n* = 5) did not result in a significant increase in IFNγ-secreting T lymphocytes as compared to a single low dose of 250 PFU of YF-EBO (*n =* 14; *p* > 0.9999 Kruskal-Wallis-test). This antiviral T_h_1-polarized CMI response was further confirmed by detection of IFNγ-secreting (cytotoxic) CD8^+^ T lymphocytes by means of flow cytometry in YF-EBO vaccinated mice after EBOV-GP peptide mix stimulation (**Fig. 3D**).

To evaluate the EBOV-specific protective efficacy induced by YF-EBO, remaining vaccinated *Ifnar*^*-/-*^ mice were infected four to six weeks post-vaccination intraperitoneally with 100 PFU of rVSV-EBOV as established before for vigorous challenge (**Fig. 3A**). Most of the sham-(*n* = 11 of 12) and all YF17D-vaccinated mice (*n*=10 of 10) rapidly developed serious disease symptoms as listed above, resulting in the development of high pain scores (**Fig. 3E**) and reaching humane endpoints as early as 2 days post-infection (**Fig. 3F**). By contrast, all YF-EBO-vaccinated mice (*n* = 11 of 11), experienced only an initial transient weight loss, associated with a moderate increase in pain scores, after which they all gained weight and remained healthy until the end of the study. A subset of mice (YF-EBO n = 3; YF17D n = 4; sham n = 3) were euthanized 3 days post-challenge to determine the infectious viral loads in different organs (liver, spleen, kidney, brain and lung). High infectious viral loads were detected in all the collected organs of sham- and YF17D-vaccinated mice (except one brain sample of a YF17D-vaccinated mouse). On the other hand, only low levels of infectious virus were detectable in the majority of the collected organs (except in the spleen) of YF-EBO-vaccinated mice on day 3 post-challenge, and no infectious virus whatsoever by the end of the study (**Fig. 3G**). Serum samples from a subset of mice that survived challenge were also analyzed. At 14 days post-challenge, while most of the control mice had already succumbed to challenge, all analyzed YF-EBO-vaccinated mice (*n* = 6 of 6) seroconverted to readily detectable levels of EBOV-specific nAb titers (**Fig. S2A**) and had markedly higher levels of EBOV-GP-specific binding antibodies (*p* = 0,0313, Wilcoxon matched-pairs signed rank test) (**Fig. S2B**) reflecting an anamnestic response following challenge.

### YFV-specific immunogenicity and protection against lethal intracranial YF17D challenge in mice

To characterize the YFV-specific immunity induced by YF-EBO, we vaccinated six-to eight-week-old *Ifnar*^*-/-*^ mice with the same single low dose of 250 PFU of YF-EBO, YF17D as a matched placebo or sham as a negative control (**Fig. 4A**).

**Figure 4:**
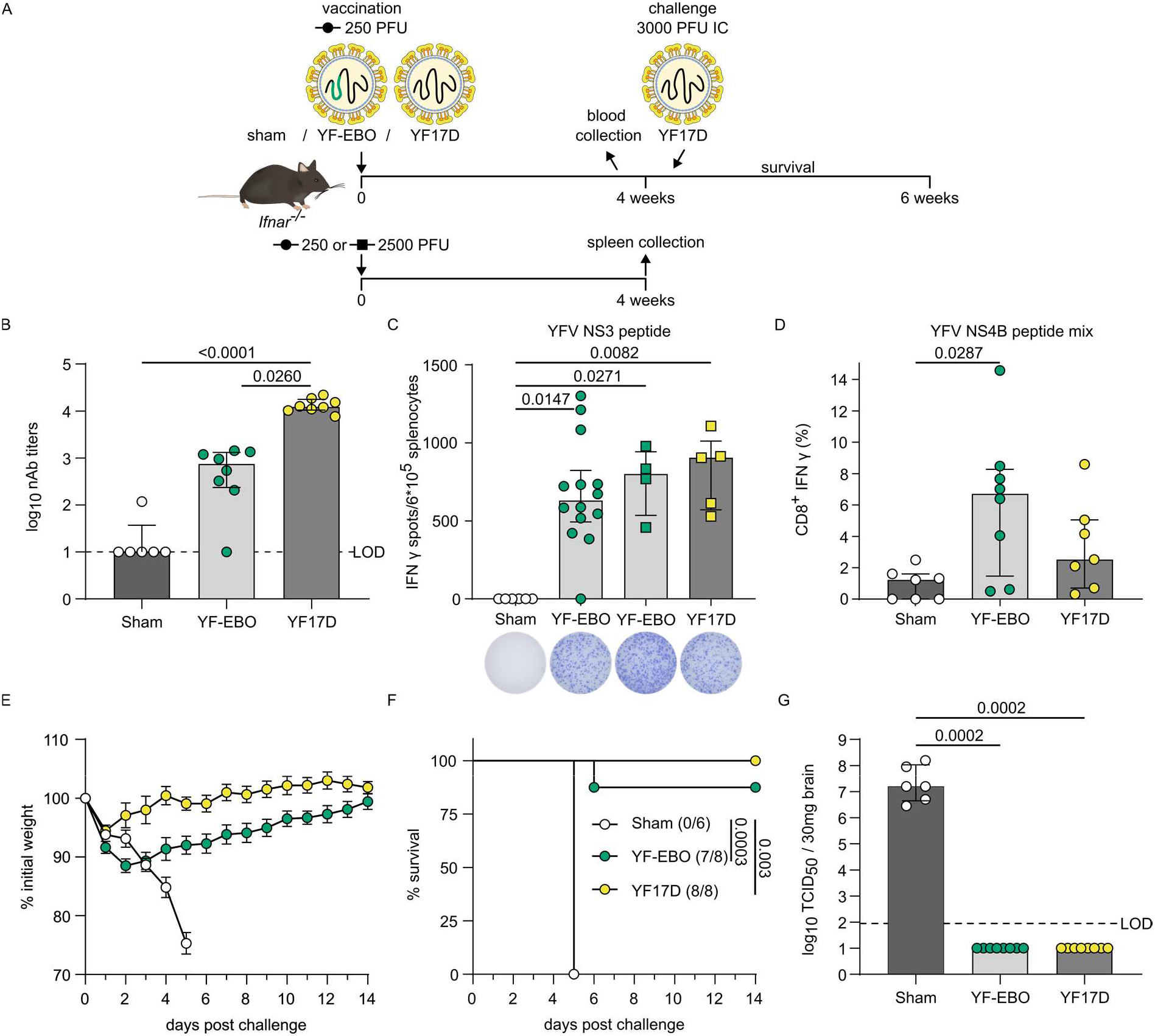
YFV-specific immunity and protection in mice. **A**. *Ifnar* ^*-/-*^ mice were vaccinated intraperitoneally with 250 (circles) or 2500 PFU (squares) of either YF-EBO, YF17D or sham. A subset of mice vaccinated with 250 PFU and all mice vaccinated with 2500 PFU were sacrificed four weeks post-vaccination for spleen collection. Remaining mice vaccinated with 250 PFU were challenged intracranially with 3000 PFU of YF17D after which they were monitored daily for two weeks for the development of disease symptoms. **B**. Pre-challenge YFV-specific neutralizing antibody (nAb) titers at four weeks post-vaccination with 250 PFU (YF-EBO *n* = 8; YF17D *n* = 8; sham *n* = 6). **C**. ELISpot counts of IFNγ-secreting cells after YFV NS3 peptide stimulation of isolated splenocytes from *Ifnar* ^*-/-*^ mice that were vaccinated with 250 PFU (circles; YF-EBO *n* = 14; sham *n* = 6) or 2500 PFU (squares; YF-EBO *n* = 5; YF17D *n* = 5). A representative image of an ELISpot well from each group is shown below the x-axis. **D**. Percentage of IFNγ-expressing CD8^+^ cells after YFV NS4B peptide pool stimulation of splenocytes from mice vaccinated with 250 PFU (YF-EBO *n* = 8; YF17D *n* = 7; sham *n* = 7). One sham mice with high background staining in ICS was excluded from the analysis because it was detected as an outlier with ROUT method (Q = 0.1%). **E**. Mean weight evolution, error bars indicate SEM (YF-EBO *n* = 8; YF17D *n* = 8; sham *n* = 6), and **F**. survival curves after challenge, number of surviving mice at study endpoint are indicated within parentheses. **G**. YF17D infectious viral loads in the brain at day of euthanasia determined by virus endpoint titration on BHK-21J cells. Dashed line indicates limit of detection (LOD). Data are median ±IQR and two-tailed Kruskal–Wallis test was applied followed by Dunn’s multiple comparison (B-D and G) and a log-rank test was applied to compare survival curves (F), significant *p*-values < 0.05 are indicated.

Four weeks post-vaccination most of the YF-EBO-vaccinated mice analyzed (*n* = 7 of 8) had seroconverted to YFV-specific nAb titers (**Fig. 4B**) although to a somewhat lower level (log_10_-transformed geometric mean titers of 2.5, 95% confidence interval of 1.8–3.4) as compared to mice vaccinated with original YF17D (*n* = 8 of 8; log_10_-transformed geometric mean titers of 4.1, 95% confidence interval of 4–4.2; *p* =0.0260 Kruskal-Wallis-test).

Virus-specific CMI responses were analyzed as before in a subset of vaccinated mice four weeks post-vaccination, yet using YFV-derived peptides as the recall antigen. Splenocytes of YF-EBO (*n* = 14) and YF17D-vaccinated mice (*n* = 5) contained significantly higher amounts of T-cells secreting IFNγ after stimulation with a MHC-I restricted YFV NS3 peptide^23^ as compared to sham-vaccinated mice (*n= 6; p* = 0,0147 and 0,0271, respectively, Kruskal-Wallis-test). A similar level of YFV-specific antiviral T_h_1-response was observed in mice when comparing both a single low dose of 250 PFU (*n* = 14) or 2500 PFU of YF-EBO (*n* = 5) with a single dose of 2500 PFU of YF17D (*n* = 4) (**Fig. 4C**). This YFV-specific antiviral T_h_1-response was further confirmed by detection of IFNγ-secreting (cytotoxic) CD8^+^ T lymphocytes by means of flow cytometry in YF-EBO vaccinated mice after YFV NS4B peptide pool stimulation (**Fig. 4D**).

To evaluate the protective efficacy induced by YF-EBO against YFV infection, remaining vaccinated *Ifnar*^*-/-*^ mice were challenged with a lethal intracranial dose of 3000 PFU of YF17D (**Fig. 4A**). Intracranial challenge has been described previously as a stringent model to evaluate protection conferred by YF17D and YF17D-vectored vaccines in mice (Ref. ^19^ and references therein). All sham-vaccinated mice (*n* = 6 of 6) developed serious disease symptoms with neurological complications such as acute weight loss, ruffled fur, hunched posture and hind limb paralysis, uniformly reaching humane endpoints as early as five days post-challenge. At contrary, the majority of YF-EBO-(*n* = 7 of 8) and all YF17D-vaccinated mice (*n* = 8 of 8) survived challenge, recovered rapidly, gained weight and did not develop any signs of disease (**Fig 4E and F**). High infectious viral loads (**Fig. 4G**) at the day of euthanasia were detected in the brains of all sham-vaccinated mice (with a median infectious viral load of 10^7.2^ TCID50/30mg brain). In contrast, no infectious virus was detected in the brains of any of the YF-EBO-vaccinated mice at the day of euthanasia; comparable to YF17D vaccination. Notably, the single YF-EBO vaccinated animal dying after intracranial challenge had detectable levels of nAb at time of infection; suggesting premature death in this case was rather linked to the harsh experimental procedure. Likewise, absence of challenge virus in the brain suggests a successful YFV immunization in all YF-EBO-vaccinated animals.

### Distinct *Ebolavirus* vaccine candidates and humoral cross-reactivity in mice

Besides EBOV, Sudan virus (SUDV), Bundibugyo virus (BDBV) and Taï forest virus (TAFV) are other highly pathogenic viruses belonging to the *Ebolavirus* genus known to cause disease in humans^1^. Therefore, in addition to YF-EBO we have generated YF17D-vectored vaccine constructs expressing their respective GP, tentatively named: YF-SUD, YF-BDB and YF-TAF (**Fig. 5A**). Comparable to YF-EBO these vaccine constructs have a smaller plaque phenotype than parental YF17D. To evaluate *Ebolavirus* cross-reactive humoral immune responses induced by these individual vaccine candidates, we hyperimmunized six-to eight-week-old *Ifnar*^*-/-*^ mice repeatedly with YF-EBO, YF-SUD, YF-BDB, or YF-TAF, and YF17D as a matched negative control. The presence of specific IgG bAb against five different *Ebolavirus*-GPs (EBOV-GP; SUDV-GP; BDBV-GP; TAFV-GP; and additionally Reston virus GP, RESTV-GP) was determined in pooled-sera of mice vaccinated 3-times with either vaccine candidate and represented in a heatmap as log_10_-transformed mean binding antibody titers (**Fig. 5B**). All vaccine candidates conferred the highest humoral immune response against their homologous target antigen. Despite the limited (55% to 73%) amino acid sequence similarity between the different *Ebolavirus* GPs, a varying degree of cross-reactive antibodies induced by the different vaccine candidates was observed, with YF-BDB inducing cross-reactive antibodies with most pronounced breadth across heterologous *Ebolavirus* species.

**Figure 5:**
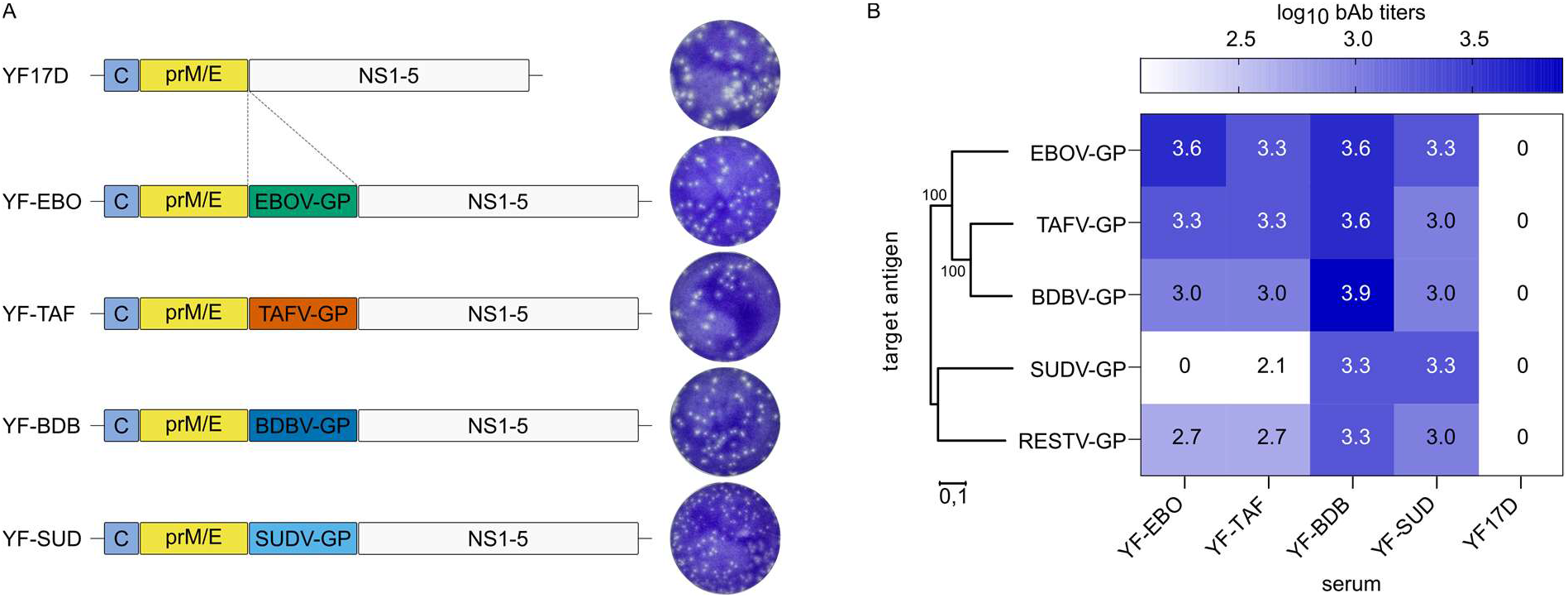
Antibody (cross)-reactivity of distinct YF17D-vectored *Ebolavirus* vaccine candidates. **A**. Schematic of YF17D-vectored *Ebolavirus* vaccine constructs and representative images of their plaque phenotypes on BHK-21J cells, fixed six days post-infection. **B**. Heatmap representing the different *Ebolavirus* GPs (EBOV-GP, SUDV-GP, TAFV-GP, BDBV-GP and RESTV-GP) log_10_-transformed mean binding antibody titers present in serum pools of *Ifnar* ^*-/-*^ mice that were hyperimmunized with either YF-EBO, YF-SUD, YF-TAF, YF-BDB or YF17D determined by IIFA (*n* = 2). Phylogenetic tree based on the amino acid sequences of the different *Ebolavirus* GPs was generated using the Neighbor-Joining method with 1000 bootstrap replications in MEGA11^51^. Small numbers at the nodes and the scale bar indicate bootstrap values and the number of amino acid substitutions per site, respectively.

## Discussion

We describe the development, preclinical safety assessment, immunogenicity and efficacy of a dual-target YF-EBO vaccine candidate in mice. Demonstration of safety and attenuation is particularly important for live-attenuated vaccines such as YF17D-vectored vaccines. YF17D that has already been used for decades, has a favorable safety profile^24^ and is therefore recommended by the World Health Organization (WHO) for all people above 9 months of age living in YFV-endemic regions, with exception of severely immunocompromised people^25^. However, there are concerns about the very rare occurrence of serious adverse events associated with YF17D vaccination (e.g. vaccine-associated neurotropic and viscerotropic disease)^26,27^. Consistent with the moderate replicative trade-off posed by the insertion of foreign genes into the YF17D backbone, we observed that YF-EBO has an attenuated phenotype *in vitro* and an improved safety profile *in vivo* as compared to YF17D, a finding previously also corroborated for a YF17D-vectored vaccine candidate against SARS-CoV-2 following a similar vaccine design ^18,28^. YF-EBO lacks the neuroinvasive properties of YF17D in immunocompromised and thus highly susceptible AG129 mice (Fig. 2B), and is attenuated compared to YF17D in a neuropotency test in neonatal BALB/c mice, while the remaining lethality induced by YF-EBO in latter model suggest sufficient potency as live-attenuated vaccine (Fig. 2C).

A single vaccine dose as low as 250 PFU of YF-EBO induced potent EBOV-GP-specific binding antibodies and EBOV-specific antiviral cellular immune response, resulting in complete protection of mice against lethal disease in a stringent surrogate EBOV infection model. Notably, the challenge model introduced and validated here uses rVSV-EBOV for vigorous challenge in immunocompromised *Ifnar*^*-/-*^ mice, allowing a first efficacy assessment under reduced BSL-2 containment. Furthermore, these data emphasize the improved safety of our YF17D-based vaccine in *Ifnar*^*-/-*^ mice (Fig. 2A) as compared to VSV-based EBOV vaccine causing lethal infection in these mice (Fig. S1). A recent study also reported that rVSV-EBOV is neurotropic in immunocompetent neonatal mice^29^, opposite to previously reported lack of neurotropism of recombinant VSV-based vaccines such as approved Ervebo® ^30^.

Immune correlates of protection for EBOV vaccines may not be universal and can vary across different vaccine platforms^31^. Yet, the presence of EBOV-GP-specific binding antibodies was shown to correlate well with protection in non-human primate models across vaccine platforms^8,32^ and may also be crucial for the observed protection in YF-EBO-vaccinated mice. Consistent with other EBOV vaccines, our vaccine candidate confers protection without inducing detectable levels of EBOV-GP-specific nAbs prior to challenge^21,22^.

Throughout the study we were limited to use immunocompromised *Ifnar*^*-/-*^ mice, since the replication of YF17D-vectored vaccines and the infection with rVSV-EBOV are highly restricted by type I IFN responses in immunocompetent wild-type mice^19,33,34^. Therefore, future mechanistic studies should address the relative contribution of humoral and cellular immune responses to the observed protection against EBOV infection in immunocompetent animal models; with extra focus on persistence of biomarkers for immunity and duration of protection.

In addition to the EBOV-specific immunogenicity, a single low dose of YF-EBO also elicited YFV-specific nAbs and T_h_1 cellular immune responses, resulting in protection against stringent intracranial YF17D challenge in *Ifnar*^*-/-*^ mice^19^. Hence, YF-EBO could also be considered as a potent YFV vaccine candidate. In this way a single dual-target vaccine could help prevent the risk of new flare-ups of EVD and concomitantly contribute to initiatives such as the Eliminate Yellow Fever Epidemics (EYE) program from the WHO^35^ aiming to increase population-wide YFV coverage in endemic regions. Likewise, latent EBOV infection persisting in EVD survivors^36–39^ could be the source of new outbreaks, meaning more frequently and in absence of otherwise rare zoonotic spill-over events ^40^. Such a dramatic change in the ecology of EVD asks for routine immunization in EBOV-endemic countries^10,11^ requiring safe and potent second-generation vaccines that can be readily deployed, ideally by single dosing as demonstrated here in principle for YF-EBO.

One of the concerns of viral vector vaccines is the impact of pre-existing anti-vector immunity on vaccine efficacy. However, we previously reported that pre-existing YFV-specific immunity in mice and hamsters did not reduce the efficacy of a similar YF17D-vectored SARS-CoV-2 vaccine candidate against SARS-CoV-2^41^. This suggests that it might also be feasible to use YF-EBO in YFV-seropositive populations in Africa.

Finally, we showed that YF17D as a vaccine vector is amenable to target different species from the *Ebolavirus* genus and can be used to develop filovirus vaccines that may induce cross-reactive binding antibodies. Future studies would have to address whether this observed cross-reactive humoral immunity is sufficient to also provide cross-protection against heterologous challenge.

In conclusion, we have developed and characterized a novel dual-target single-shot YF17D-vectored EBOV vaccine candidate that provides protection against both EBOV and YFV infection in mice, which warrants further (pre-)clinical development.

## Materials and methods

### Cells and viruses

BHK-21J (baby hamster kidney fibroblasts) cells were provided by P. Bredenbeek and maintained in minimum essential medium (MEM) (Gibco), Vero E6 (African green monkey kidney, ATCC CRL-1586) and HEK293T (human embryonic kidney cells, ATCC CRL-3216) cells were maintained in Dulbecco’s modified Eagle medium (DMEM) (Gibco). All media were supplemented with 10% fetal bovine serum (Hyclone), 2 mM L-glutamine (Gibco), 1% sodium bicarbonate (Gibco). BSR-T7/5 (T7 RNA polymerase expressing BHK-21) cells were provided by I. Goodfellow and kept in DMEM supplemented with 0.5 mg/ml geneticin (Gibco).

Replication-competent recombinant VSV-EBOV was generated by cloning the Makona EBOV-GP sequence (GenBank: KM233070.1) into the multiple cloning site of pVSV-ΔG-GFP-2.6, kindly provided by M. A. Whitt, using MluI and XmaI restriction sites. Viral recovery of rVSV-EBOV has been performed as previously described^42,43^ by co-transfection of BSR-T7/5 cells with 2.5 µg of pVSV-ΔG-EBOV-GP-GFP-2.6 and VSV helper plasmids encoding for VSV-N, P, L, and G proteins with a ratio of 0.87:1.43:2.25:0.5 µg of each plasmid, respectively, and amplification of the rescued virus on Vero E6 cells. This first recovery virus was plaque-purified and amplified on Vero E6 cells to generate a homogeneous virus stock of rVSV-EBOV. Sanger sequencing and immunofluorescent staining of infected cells confirmed the presence of the EBOV-GP sequence in generated virus stocks. Virus titers were determined by plaque assay on Vero E6 cells, expressed as plaque forming units (PFU)/mL as described^44^.

YF17D virus, Stamaril® (lot G5400) (Sanofi-Pasteur) used for intracranial challenge experiments was passaged three times in Vero E6 cells before use. Virus titers were determined by plaque assay on BHK-21J cells as described^44^.

### Vaccine design and construction

Vaccine constructs were generated using an infectious cDNA clone of YF17D (in an inducible BAC expression vector pShuttle-YF17D)^44–46^ using standard molecular biology techniques essentially as described^18^. In brief, YF-EBO was engineered by inserting the sequence of EBOV-GP strain Makona (GenBank: KM233070.1; corresponding to aa 33-676, i.e. excluding the EBOV-GP signal peptide sequence; obtained after PCR on overlapping synthetic cDNA fragments; IDT) into the full-length genome of YF17D (GenBank: X03700) as translational in-frame fusion within the YF-E/NS1 intergenic region^17,47^, followed by transmembrane domains derived from WNV to ensure a proper YF topology and correct ER luminal expression of the GP antigens in the YF backbone^18^. Infectious recombinant viruses were generated by transfection into BHK-21J cells and harvesting supernatants harvested three or four days post-transfection when most of the cells showed signs of cytopathic effect (CPE). Infectious virus titers (PFU/ml) were determined by a plaque assay as previously described^44^. The presence of inserted sequences in thus generated vaccine virus stocks was confirmed by Sanger sequencing. YF-SUD expressing SUDV-GP (GenBank: MH121163.1), YF-TAF expressing TAFV-GP (GenBank: NC_014372.1), and YF-BDB expressing BDBV-GP (GenBank: KU182911.1) were generated and rescued accordingly.

### Virus growth kinetics

Viral growth kinetics were determined by infection of BHK-21J cells at an MOI of 0.01. One hour post-infection, cells were washed and medium was replaced. Supernatant samples were collected every day for 5 days and frozen at -80°C. Viral yields were determined by virus endpoint titration on BHK-21J cells.

### Immunofluorescent staining

In vitro antigen expression of the vaccine candidate was verified by immunofluorescent staining as previously described^44^. In brief, BHK-21J cells were infected with the YF-EBO vaccine candidate. Infected cells were stained two days after infection. For detection of yellow fever virus antigens, polyclonal hamster anti-YF17D antiserum was used. For detection of EBOV-GP antigen, mouse anti-Zaire Ebola virus GP monoclonal antibody 4F3 (IBT Bioservices, 1:500 dilution) was used. Secondary antibodies were goat anti-mouse Alexa Fluor 594 (Life Technologies; 1:500 dilution) and rabbit anti-hamster FITC (Jackson Immuno Research; 1:500 dilution). Cells were counterstained with DAPI (Sigma). All confocal fluorescent images were acquired using the same settings on a Leica TCS SP5 confocal microscope, using a HCX PL APO 63× (NA 1.2) water immersion objective.

### Animal experiments

Six-to twelve-week-old male and female *Ifnar*^*-/-*^ mice (type-I-interferon-receptor-deficient C57BL/6 mice) and six-to eight-week-old male and female AG129 mice (type-I-and-II-interferon-receptor-deficient 129 mice) were used throughout this study and were bred in-house. Five days old BALB/c pups were purchased from Janvier Laboratories. Animals were kept in individually ventilated filtertop cages (Sealsafe Plus, Tecniplast) with a maximum of 5 mice per cage. Mice were provided with food and water ad libitum and cage enrichment (cotton and cardboard play tunnel/shelter). Mice were randomly assigned to different experimental treatment groups within one cage, to avoid any cage effect. Projects were approved by the KU Leuven ethical committee (P030-2021, P140-2016 and P100-2019), following institutional guidelines approved by the Federation of European Laboratory Animal Science Associations (FELASA). Animal health was monitored throughout the study and animals were euthanized at experimental endpoint or when pre-defined humane endpoints were reached by intraperitoneal administration of 100µl of Dolethal® (200mg/ml sodium pentobarbital, Vétoquinol SA, Magny-Vernois, France).

To evaluate neurovirulence and neurotropism, BALB/c mice pups and AG129 mice were, respectively, intracranially or intraperitoneally inoculated with the indicated amount of PFU of YF17D or YF-EBO and monitored daily for morbidity and mortality after inoculation.

To determine rVSV-EBOV infectivity, ten-to twelve-week-old *Ifnar*^*-/-*^ mice were infected intraperitoneally with either 100 or 100.000 PFU of rVSV-EBOV and monitored daily for two weeks for the development of disease using the Institutional Animal Care and Use Committee (IACUC) pain-scoring list (supplementary table S1) including following parameters: body weight changes, body-condition score, physical appearance and behavior. Mice were euthanized when they reached a total score of three or more or when they developed one of the following symptoms: paresis, paralysis or a hunched back. Organs (kidney, liver, spleen, brain and lung) were collected at the day of euthanasia for endpoint virus titrations.

Six-to eight-week-old *Ifnar*^*-/-*^ mice were intraperitoneally vaccinated with 250 or 2500 PFU of YF-EBO as indicated. As a control, two groups were vaccinated with either YF17D or sham (MEM with 2% FBS). Of note, the intraperitoneal route of immunization for YF17D and derivatives thereof in small animal models has been intensively validated by us before^18,19,41,48^ and successfully bridged to the clinically relevant subcutaneous route in a non-human primate model^18^. All mice were bled four weeks post-vaccination for indirect immunofluorescence assays (IIFA) and serum neutralization tests (SNT). A subset of mice was euthanized at four weeks post-vaccination and spleens were collected for ELISpot cytokine detection or flow cytometry analysis. For rVSV-EBOV challenge, mice were intraperitoneally infected with a lethal dose of 100 PFU of rVSV-EBOV four to six weeks post-vaccination. Mice were monitored daily for two weeks for the development of disease using the IACUC pain-scoring list (supplementary table S1) and were euthanized when humane endpoints were reached, as described above. A subset of mice was euthanized at 3 days post-challenge and organs (kidney, liver, spleen, brain and lung) were collected for endpoint virus titration. For YFV challenge, mice were inoculated with a lethal dose of YF17D via the intracranial route, as previously described^19^. Briefly, four weeks after intraperitoneal vaccination with 250 PFU of YF-EBO, YF17D, or sham, mice were anaesthetized by intraperitoneal injection of a mixture of xylazine (16 mg kg−1, XYL-M, V.M.D.), ketamine (40 mg kg−1, Nimatek, EuroVet) and atropine (0.2 mg kg−1, Sterop). After confirmed anesthesia, mice were inoculated intracranially with 30 μl containing 3000 PFU of YF17D, and then monitored daily for signs of disease and weight change for two weeks. Mice were euthanized on the basis of morbidity (paralysis, paresis, hunched posture and ruffled fur) or weight loss of more than 20% or a quick weight loss of more than 10% in 1-2 days. Brains were collected at the day of euthanasia for endpoint virus titration.

Hyperimmune serum was generated by vaccinating *Ifnar*^*-/-*^ mice three times intraperitoneally in three consecutive weeks with the respective vaccine candidate (YF-EBO, YF-TAF, YF-BDB, YF-SUD, or YF17D). Serum was collected 5 weeks after the first vaccination and pooled for each individual vaccine candidate for analysis in an IIFA for the presence of binding antibodies against distinct *Ebolavirus* GPs.

### Detection of total binding IgG by indirect immunofluorescence assay (IIFA)

To detect EBOV-specific binding antibodies in mouse serum, an in-house-developed IIFA was used as described^18^. Briefly, HEK293T cells seeded in 96-well plates were transiently transfected with either pCMV-EBOV-GP-IRES-RFP or pCMV-RABV-CVS-G-IRES-RFP causing overexpression of the target antigen (EBOV-GP) and RFP or an irrelevant glycoprotein (RABV-CVS-G) and RFP, respectively. To determine EBOV-GP binding antibody end titers, 1/2 serial serum dilutions were made in parallel on EBOV-GP-expressing cells and RABV-CVS-G expressing cells. Goat-anti-mouse IgG Alexa Fluor 647 (A-21236, Life Technologies; 1:500 dilution) were used as secondary antibody. After counterstaining with DAPI, fluorescence in the red channel (excitation at 560 nm) and the far-red channel (excitation at 650 nm) was measured with a Cell Insight CX5 High Content Screening platform (Thermo Fischer Scientific). Specific EBOV-GP staining is characterized by cytoplasmic (endoplasmic reticulum) enrichment in the far-red channel. To quantify this specific EBOV-GP staining, the difference in cytoplasmic and nuclear signal for the RABV-CVS-G expressing cells was subtracted from the difference in cytoplasmic and nuclear signal for the EBOV-GP expressing cells. All positive values were considered as specific EBOV-GP staining. The IIFA end titer of a sample is defined as the highest dilution that scored positive in this way. Cross-reactive binding antibodies against distinct *Ebolavirus* GPs in hyperimmune serum were determined in a similar way with an IIFA on transfected cells that expressed the respective target antigen (EBOV-GP, TAFV-GP, BDBV-GP, SUDV-GP or RESTV-GP) and RFP, the read-out was done on a DMi8 microscope (Leica) with fixed microscope settings.

### YFV seroneutralization test

To quantify YFV-specific neutralizing antibodies, serum samples were tested in a SNT assay using an mCherry-tagged YF17D reporter virus as described in great detail^49^.

### rVSV-EBOV seroneutralization test

To quantify EBOV neutralizing antibodies, serum samples were serially diluted and incubated for 1 h at 37 °C with 1000 PFU of rVSV-EBOV, which contains an additional transcription unit encoding for GFP, and inoculated overnight on Vero E6 cells. The percentage of GFP expressing cells was quantified on a Cell Insight CX5 High Content Screening platform (Thermo Fischer Scientific) and half-maximal inhibitory serum neutralization titers (SNT_50_) values were determined, as described^50^. Data were normalized to a virus (100%) and cell control (0%) and fitted to a nonlinear regression curve to determine SNT_50_ values in Graphpad Prism (GraphPad Software).

### ELISpot

Collected spleens were passed through a cell-strainer (70 μm) to obtain single-cell suspensions and red blood cells were lysed using RBC lysis-buffer (eBioscience). ELISpot assays for the detection of IFNγ-secreting mouse splenocytes were performed with mouse IFNγ kit (ImmunoSpot MIFNG-1M/5, CTL Europe) according to manufacturer instructions. Briefly, 6×10^5^ Splenocytes were stimulated with either an EBOV-GP peptide pool (PepMix™ Zaire Ebola (GP/Kikwit-95), 0.25µM/peptide, JPT peptide technologies), a YF17D NS3 peptide (ATLTYRML, NS3_268–275_, 5 μM, Eurogentec, Seraing, Belgium) or an ovalbumin peptide (SIINFEKL, OVA_257-264_, 5 μM, Eurogentec, Seraing, Belgium). After 48 h of incubation at 37°C, IFNγ spots were visualized by stepwise addition of a biotinylated detection antibody, a streptavidin-enzyme conjugate and the substrate. Spots were counted using an ImmunoSpot S6 Universal Reader (CTL Europe) and normalized by subtracting spots numbers from control samples (incubated with irrelevant ovalbumin peptide) from the spot numbers of corresponding stimulated samples. Negative values were corrected to zero.

### ICS and flow cytometry

Fresh mouse splenocytes were stimulated with 1.2 µM/peptide of EBOV-GP peptide pool (PepMix™ Zaire Ebola (GP/Kikwit-95) JPT peptide technologies), or 1.5 µM/peptide YF17D NS4B peptide pool (PepMix™ Yellow fever (NS4B) JPT peptide technologies) or with medium supplemented with DMSO. After 48 h of incubation at 37°C, cells were treated with 1x Brefeldin A (ThermoFisher Scientific) for 4 h at 37°C, thoroughly washed and stained with Zombie Aqua (Zombie Aqua™ Fixable Viability Kit, Biolegend) and FcgR block (0.5 µl/well, FcR Blocking Reagent mouse, Miltenyi Biotec) for 15 min at RT in the dark. Cells were washed again and were stained with surface markers BUV395 anti-CD3 (1:167 dilution, clone 17A2, ThermoFisher Scientific), BV785 anti-CD4 (1:100 dilution, clone GK1.5, Biolegend), and APC/Cyanine7 anti-CD8 (1:100 dilution, clone 53-6.7, Biolegend) in Brilliant Staining Buffer (BD Biosciences) and incubated for 25 min on ice in the dark. The cells were thoroughly washed again followed by fixation and permeabilization (eBioscience™ Foxp3/Transcription Factor Staining Buffer Set, Invitrogen) for 30 min on ice in the dark. Subsequently the cells were washed with permeabilization buffer (eBioscience™ Foxp3 / Transcription Factor Staining Buffer Set, Invitrogen) and stained for 45 min on ice in the dark with an APC anti-IFNγ (1:100 dilution, clone XMG1.2, Biolegend) intracellular marker. After a final washing step, analysis was performed with a BD LSRFortessa X20 device. Data was gated using FlowJo software 10.8.1 (BD) (strategy shown in supplementary Fig. S3) and all samples were normalized by subtraction of peptide-stimulated cells by background from non-stimulated cells of the same animal.

### Endpoint virus titrations

To quantify infectious rVSV-EBOV or YF17D viral loads in the organs of challenged mice, endpoint titrations were performed on confluent Vero E6 cells or BHK21J cells, respectively. A piece of each collected organ (kidney, liver, spleen, brain and lung) weighing approximately 30 mg was homogenized using bead disruption (Precellys) in 350µl MEM and centrifuged (10.000 rpm,10min, 4°C) to pellet cell debris. Viral loads were calculated by the Reed and Muench method and expressed as TCID50 per 30mg tissue.

### Data analysis

All statistical analyses were performed using GraphPad Prism 9 software (GraphPad, San Diego, CA, USA). Results are presented as medians ± IQR or mean ± SEM as indicated. Statistical differences for multiple comparisons were analyzed using Kruskal–Wallis with Dunn’s multiple comparisons test, survival curves were analyzed using a log-rank test (Mantel-Cox), repeated measures were analyzed using a Wilcoxon matched-pairs signed rank test and all tests were considered statistically significant at *p* values <0.05.

## Acknowledgements

We are grateful to Prof. Michael A. Whitt (University of Tennessee Health Science Center) for generously sharing VSV plasmids. We thank Katrien Geerts, Jasper Rymenants, Carolien De Keyzer and Els Brouwers (PharmAbs, KU Leuven) for excellent technical assistance with cell culture, plaque assays and animal work; Robbert Boudewijns for sharing ELISpot protocol; Niraj Mishra and Mahadesh Prasad Arkalagud Javarappa for intracranial injections; Jasmine Paulissen and Madina Rasulova (TPVC) for diligent YFV-SNT assays and ELISpot-plate read-outs; Els Vanstreels for confocal microscopy set-up; Elisabeth Heylen for carefully evaluating the risk assessment; the staff of the Rega animalium for taking great care of the animals and finally Lut Overbergh (Laboratory of Clinical and Experimental Endocrinology, KU Leuven) for critically reading and correcting this manuscript.

Current work was supported by the Flemish Research Foundation (FWO) Excellence of Science (EOS) program (No. 30981113; VirEOS project and No. 40007527; VirEOS2), the European Union’s Horizon 2020 research and innovation program (No. 733176; RABYD-VAX consortium). V.L. acknowledges a research assistant fellowship from KU Leuven, J.M. grant support from the Chinese Scholarship Council (CSC; No. 201706760059) and K.D. from KU Leuven Internal Funds (C3/19/057; Lab of Excellence).

## Author contributions

Designed experiments: K.D., L.S.-F., V.L; vaccine design: L.S.-F, K.D.; molecular work: L.S.-F, S.D., V.L.; animal experiments: L.K., V.L.; serology: L.K, W.C., T.V., H.J.T., V.L.; Elispot & flow: L.K., J.M., V.L.; analyzed data: L.K., V.L.; provided advice on the interpretation of data: K.D., L.S.-F.; wrote the original draft with input from co-authors: V.L.; wrote the final draft: K.D., L.S.-F., L.K., V.L.; supervised the study: K.D., L.S.-F., J.N.; acquired funding: J.N., K.D.; all authors approved the final manuscript.

## Competing interests

K.D., L.S.F., V.L. and J.N. are mentioned as inventors on patent applications related to the discovery and use of YF17D-vectored filovirus vaccines. The other authors declare no competing interest.

## Data availability

All data supporting the findings in this study are available from the corresponding author upon request.

---

**Supplementary figure S1:**
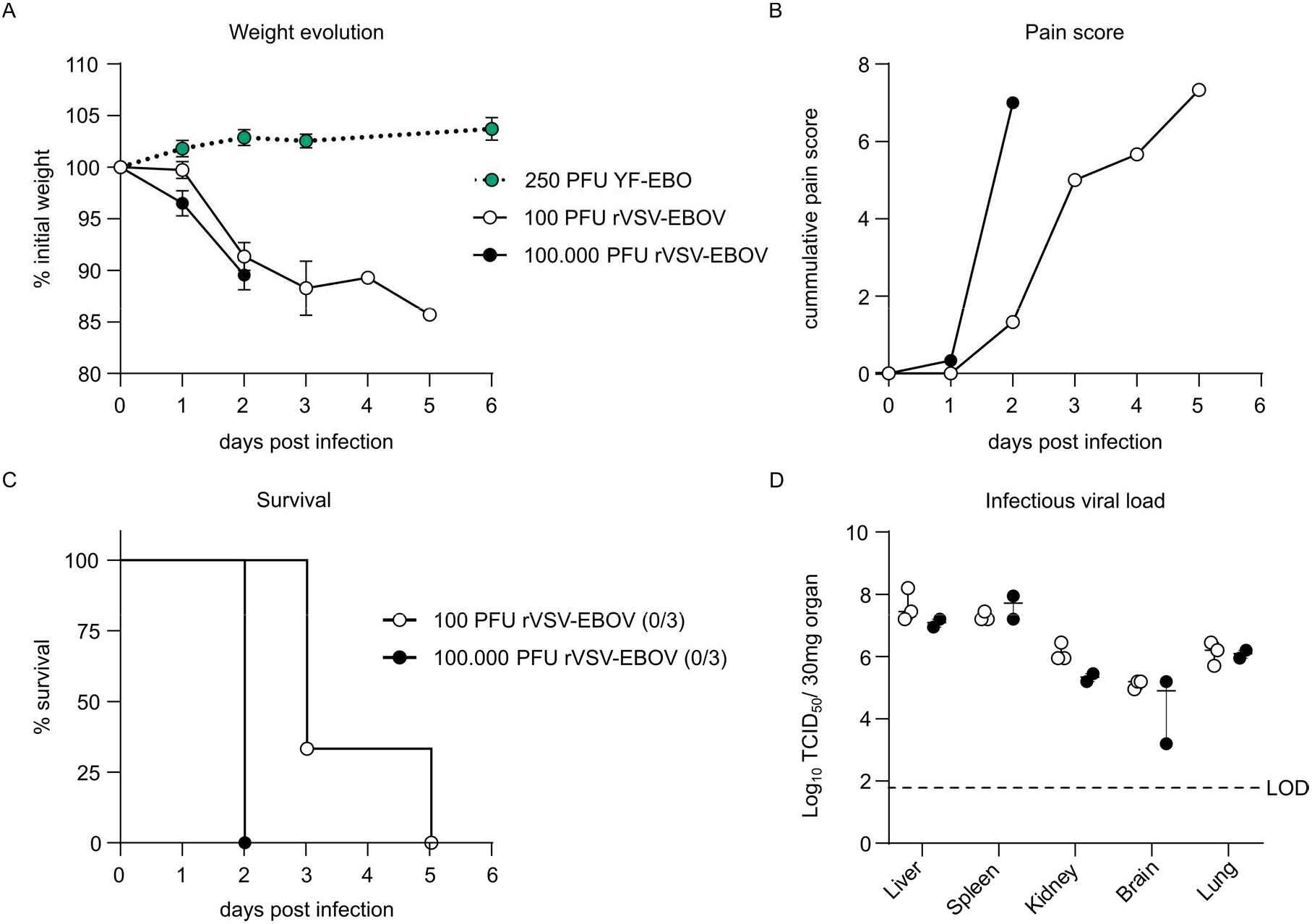
rVSV-EBOV infection in mice. *Ifnar* ^*-/-*^ mice were infected intraperitoneally with 100 (*n* = 3) or 100.000 PFU (*n* = 3) of rVSV-EBOV and monitored for the development of disease symptoms. **A**. Weight evolution after infection with rVSV-EBOV, as a comparison the dotted line represents weight evolution of *Ifnar*^*-/-*^ mice (*n* = 6) after intraperitoneal inoculation with 250 PFU of YF-EBO (data as in Figure 2A). Error bars represent SEM. **B**. Mean cumulative pain score, based on IACUC parameters (see Supplementary table S1) including: body weight changes, body condition score, behaviour and physical appearance. **C**. Survival curve after rVSV-EBOV infection. The number of surviving mice at study endpoint are indicated within parentheses. **D**. rVSV-EBOV infectious viral loads in different organs collected at the day of euthanasia and quantified by virus titration on Vero E6 cells. Data are median ±IQR, dashed line represents limit of detection (LOD) (D).

**Supplementary figure S2:**
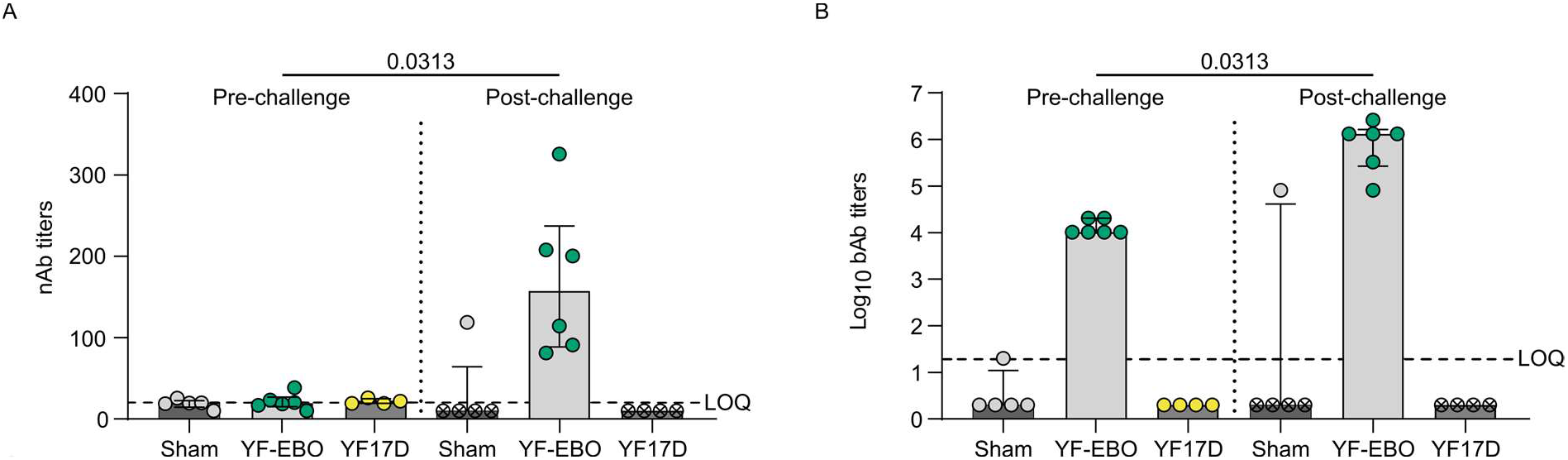
comparison of EBOV-specific humoral immunity pre- and post-challenge. Related to figure 3. Pre-challenge (four weeks post-vaccination) and post-challenge (2 weeks post-challenge) **A**. EBOV-GP-specific neutralizing antibody (nAb) titers determined by rVSV-EBOV seroneutralization test and **B**. EBOV-GP-specific IgG binding antibody (bAb) titers determined by IIFA, present in serum of mice vaccinated with 250 PFU (YF-EBO *n =* 6; YF17D *n* = 4; sham *n* = 5). Mice that already succumbed to rVSV-EBOV infection are represented with an ‘x’. Dashed line represents limit of quantification (LOQ), data are median ±IQR and two-tailed Wilcoxon matched-pairs signed rank test was applied, significant *p*-values < 0.05 are indicated (A-B).

**Supplementary figure S3:**
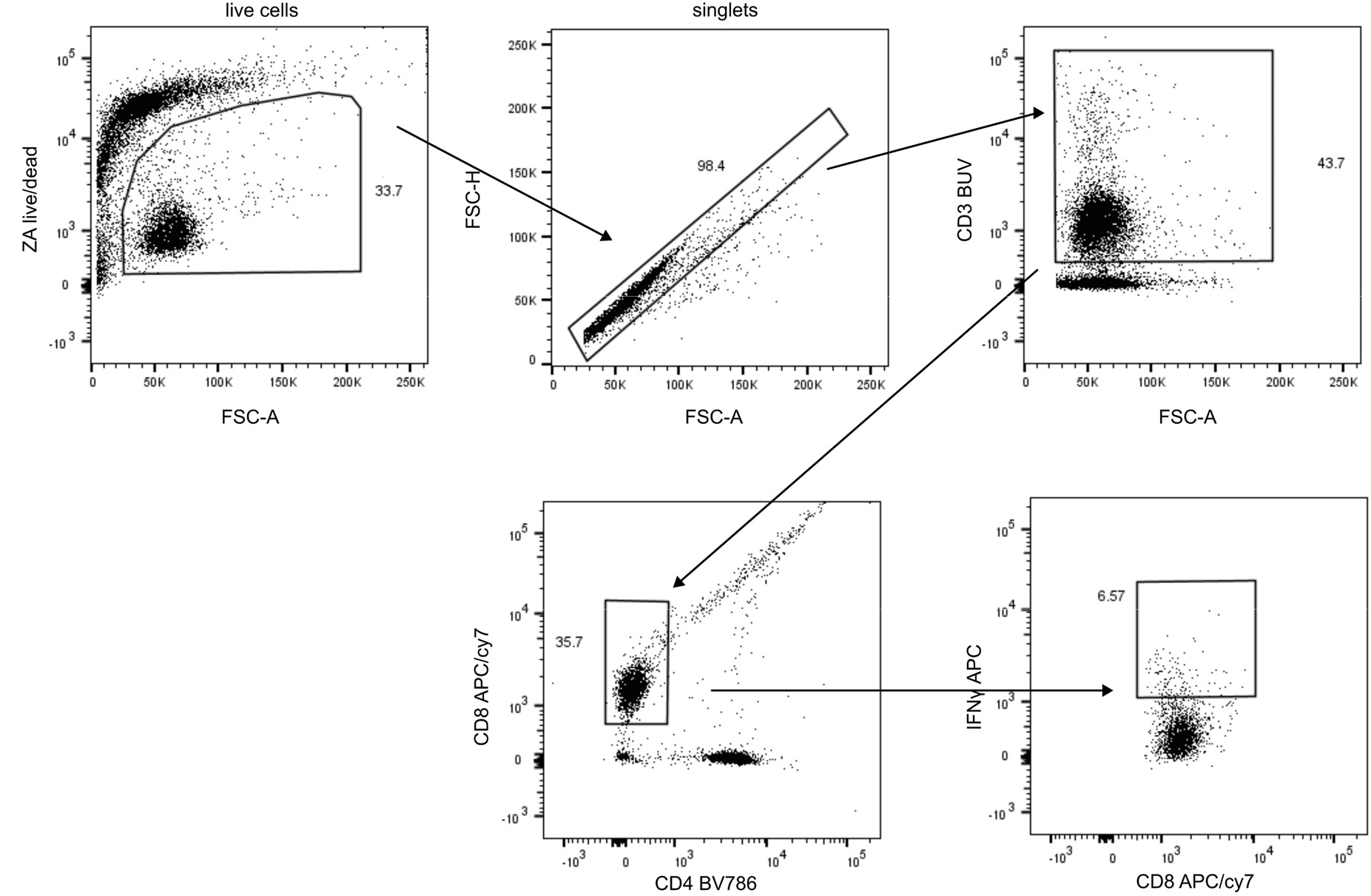
gating strategy. First, live cells were selected by gating out Zombie Aqua (ZA)-positive and low forward scatter (FSC) events. Then, doublets were eliminated in an FSC-H versus FSC-A plot. T cells (CD3+) were stratified into CD8 T cells (CD8+). Boundaries defining positive and negative populations for intracellular marker (IFNγ) were set on the basis of non-stimulated control samples.

**Supplementary table S1:** IACUC pain-scoring list.

| Body weight changes | 0 | Normal |
| --- | --- | --- |
|  | 1 | <10% Weight loss |
|  | 2 | 10-15% Weight loss |
|  | 3 | >20% Weight loss |
| Body-condition score | 0 | Well-conditioned (vertebrae, pelvic, or spinal bones not prominent) |
|  | 1 | Under-conditioned (evident vertebral segmentation; pelvic bones readily palpable) |
|  | 2 | Emaciation (skeletal structures extremely prominent; little or no flesh cover) |
| Physical appearance | 0 | Normal |
|  | 1 | Lack of grooming |
|  | 2 | Rough coat, nasal/ocular discharge |
|  | 3 | Very rough coat, abnormal posture |
| Behaviour | 0 | Normal |
|  | 1 | Minor changes (e.g., limping) |
|  | 2 | Abnormal; reduced mobility, inactive |
|  | 3 | Unsolicited vocalizations, self-mutilation, either very restless or immobile |

